# Planted origin of shade trees: a robust determinant of cocoa yield among smallholder farmers in Côte d’Ivoire

**DOI:** 10.64898/2026.09.24.754229

**Authors:** Beda Innocent Adji, Assiri Assiri Alexis, Assi Maryse Evelyne, Kassin Koffi Emmanuel, Doffou Sélastique Akaffou

**Affiliations:** Jean Lorougnon Guédé University, Agroforestry Department, BP 150 Daloa, Côte d’Ivoire; Cocoa program, National Center for Agronomic Research (CNRA), BP 808 Divo, Côte d’Ivoire

**Keywords:** planted origin of shade trees, cocoa agroforestry, West African smallholder, *LASSO*, agricultural human capital, Côte d’Ivoire

## Abstract

**CONTEXT:** While the agroeconomic literature on the determinants of cocoa yield among West African smallholders is abundant, it remains dominated by descriptive studies or work conducted at an aggregated regional scale, leaving open the question of the role of shade-tree management practices at the scale of the individual plantation.

**OBJECTIVE:** This study tests whether shade-tree management practices and the farmer’s socio-demographic profile explain variation in cocoa yield, using a nationwide sample of 409 plantations covering, for the first time, the three major Ivorian production zones (« loops ») (98, 151 and 160 plantations for loops 1, 2 and 3, respectively).

**METHODS:** A univariate screening of 27 variables, a mixed model with a random intercept by village, a production-function specification, LASSO variable selection, and a random forest were applied, all evaluated by 5-fold cross-validation.

**RESULTS AND CONCLUSIONS:** The five approaches converge on a robust result: the deliberately planted origin of shade trees (as opposed to a residual or spontaneous origin) is the strongest and most stable determinant of yield, with a mean gap of 332 versus 183 kg/ha/year. This effect withstands four successive robustness checks: it remains significant after simultaneous adjustment for age, plantation size, production zone, technical extension, and farmer education; it is not driven by a handful of extreme plantations; it holds within each of the three production zones taken separately rather than in only one of them; and, taken in isolation, it retains a positive out-of-sample predictive power (cross-validated *R*^*2*^ ≈ 0.05). A second group of robust determinants of more modest magnitude emerges for agricultural technical extension (ANADER/CNRA/SATMACI) and farmer education level; shade-tree alignment shows a signal in the same direction, consistent with recent independent work in Côte d’Ivoire, but becomes statistically marginal once adjusted for these other factors. The full multivariate model reaches a cross-validated *R*^*2*^ of around 0.07-0.08.

**SIGNIFICANCE:** This signal, undetectable in an analysis restricted to loop 1 alone (*n*=98) for lack of statistical power, confirms the value of nationwide sampling for detecting modest but real agronomic determinants, and argues for integrating shade-tree origin into agroforestry extension programmes.

## 1. Introduction

Whether, and how, shade trees associated with cocoa affect the latter’s yield is today a central question in the debate on the sustainability of West African cocoa farming, particularly since the entry into force of the European Union’s « deforestation-free » products regulation (Regulation (EU) 2023/1115; Union Européenne, 2023), which encourages exporting countries to rigorously document their agroforestry practices. The international literature has shown that shaded cocoa systems can reconcile production with biodiversity conservation, but that this outcome depends closely on the structure and management intensity of the shade canopy (Clough et al., 2011; Tscharntke et al., 2011), and that at the stand level it is the functional traits of shade trees (height, architecture, phenology), rather than their mere presence or origin, that determine the services provided (Isaac et al., 2024).

This observation is reinforced by more recent work: in West Africa, the vulnerability of cocoa agroforestry systems to climate change depends directly on shade-canopy structure (Ariza-Salamanca et al., 2025), while moderate shade can reconcile emission reductions with maintained profitability in Ghana (Hawkins et al., 2024); more broadly, the potential of agroforestry for an emissions-intensive agricultural commodity such as cocoa remains underexploited (Becker et al., 2025). On the governance side, compliance with the European Union’s « deforestation-free » products regulation in Côte d’Ivoire relies heavily on local cooperative structures, whose effective capacity to enforce these new documentary requirements remains variable (Moluh Njoya et al., 2025).

In Côte d’Ivoire, this question remains poorly documented at the scale of the individual plantation. The Master’s thesis underlying this work (Adji et al., 2016) qualitatively documented, across 474 plantations and three major cocoa-growing loops (representing the three main historical cocoa-bean production zones of Côte d’Ivoire), a synergy between the presence of shade trees and perceived yield, but without formal statistical testing and at an aggregated scale. A broader agronomic literature on the determinants of cocoa yield in West Africa identifies a set of management factors (planting material, fertilisation, technical extension, orchard age) as explaining a substantial share of the yield gaps observed between farms (Abdulai et al., 2020, for Ghana; Assiri et al., 2009, 2016, for Côte d’Ivoire). A recent quantification of this yield gap in Ghana (Asante et al., 2022) confirms that it remains considerable even among farmers following recommended best practices, while a machine-learning approach applied to Ghana and Côte d’Ivoire (Obahoundje et al., 2026) highlights the value of non-linear methods, such as those used here, for capturing complex interactions between climatic and agronomic determinants of cocoa yield. This raises the question of whether variables specifically describing shade-tree management could likewise explain part of the yield variation observed in our own dataset. A very recent and independent study, published in May 2026 in Scientific Reports on a distinct sample of Ivorian plantations, provides an important point of reference: Yéo et al. (2026) show that the alignment of planted trees (+51% yield), mineral fertilisation (+29%), and the use of certified planting material (+23%) are significant determinants of cocoa yield, while older farmer age is associated with lower productivity. This work offers a direct and recent benchmark against which to assess whether comparable determinants are detectable in our own sample.

Beyond shade-tree management practices as such, two further categories of potential cocoa-yield determinants, widely debated in the agroeconomic literature, deserve to be tested concurrently rather than treated as mere control variables: the relationship between farm size and productivity, whose sign and very existence remain the subject of a classic and unresolved debate in African agricultural economics (Ali and Deininger, 2015, for Rwanda), and the role of the farmer’s human capital (education level, access to technical extension) as a determinant of the adoption of productive practices (Alene and Manyong, 2007, pour le Nigeria). On the first point, recent re-examinations of this relationship, covering a wider range of farm sizes, reach similar conclusions in Kenya (Muyanga and Jayne, 2019) and Nigeria (Omotilewa et al., 2021), while a study focusing specifically on Ghanaian cocoa concludes that spatial and management determinants, rather than size itself, explain most of the productivity variation between farms (Bentum et al., 2026). On the second point, a recent meta-analysis confirms an overall positive, though context-dependent, effect of technical extension on agricultural productivity (Ogundari, 2022), a result corroborated in Ghana by recent panel data (Aremu et al., 2025) and complemented, for the educational component of human capital, by recent South African data (Baiyegunhi, 2024). The present article documents these two categories of determinants alongside shade-tree management practices, with the same methodological rigour.

This study addresses the following research question: do shade-tree management practices, combined with the farmer’s socio-demographic characteristics, explain the variation in individual cocoa yield across the three main production zones of Côte d’Ivoire, and do these effects withstand out-of-sample validation rather than resting solely on in-sample fit? We test three pre-specified hypotheses:

✓ H1 (deliberate planting): Plantations whose shade trees are predominantly of deliberately planted origin (rather than residual forest trees retained during clearing or uncontrolled natural regeneration) achieve higher individual cocoa yield, independently of the production zone;
✓ H2 (human capital): Access to formal technical extension (ANADER, CNRA, SATMACI, or a certification programme) and the farmer’s education level are positively associated with yield, consistent with a human-capital interpretation of yield gaps among smallholders;
✓ H3 (farm size and cooperative membership; competing/null hypotheses): Neither plantation size nor formal cooperative membership, taken in isolation, constitutes a robust determinant of yield once management-practice and human-capital covariates are accounted for; we test these two factors as genuine competing hypotheses rather than as mere control covariates, and report null results with the same transparency as positive ones.

We further put forward the methodological hypothesis that a nationwide sample covering the three production zones has the statistical power and covariate variance needed to detect management-practice determinants of modest magnitude, which a sub-sample restricted to a single zone would fail to detect. To test H1 through H3, the objective of the present paper is to formally test, on the 409 plantations from the three production loops with an individual yield value, whether shade-tree management practices and a set of the farmer’s socio-demographic covariates explain the variation in cocoa yield, by drawing on a set of complementary methods (univariate screening, mixed model, production-function specification, *LASSO* variable selection, random forest), all honestly evaluated by cross-validation rather than by in-sample fit alone, and systematically checked for consistency across the three production zones rather than in only one of them.

## 2. Materials and methods

The data used here come from the field survey documented in a Master’s thesis in Agroforestry (Adji et al., 2016; 474 plantations recorded, 103 localities, three cocoa-growing loops), using the survey forms appended to this document (Supplementary material: S1). The present paper focuses specifically on the sub-sample of plantations with a recorded individual yield, across the three loops.

### 2.1. Study area

The data come from a field survey conducted in the three main cocoa-growing zones of Côte d’Ivoire, hereafter referred to as « loops » (boucles), a grouping convention used throughout this article and inherited from the source thesis (Adji et al., 2016). The three loops span a clear north-south bioclimatic gradient (Fig. 1), from the mesophilous, pre-forest sector partly covered by loop 2 (Centre/Centre-West) to the dense-forest sector characteristic of loops 1 (East/South-East) and 3 (South/South-West), and together cover 103 survey localities spread across sixteen departments and eight administrative regions (Table 1). Loop 1 (East/South-East) comprises 4 departments (Abengourou, Adzopé, Akoupé, Agboville) spread across 3 regions and 28 survey sites; loop 2 (Centre/Centre-West) comprises 8 departments (Djékanou, Toumodi, Daloa, Vavoua, Issia, Zoukougbeu, Yamoussoukro, Bouaflé) spread across 4 regions and 51 survey sites; and loop 3 (South/South-West) comprises 4 departments (Divo, Guitry, Soubré, Tiassalé) spread across 3 regions and 24 survey sites (Table 1). This geographic arrangement underlies the production-zone (loop) covariate used systematically throughout the statistical analysis below.

**Fig. 1.**
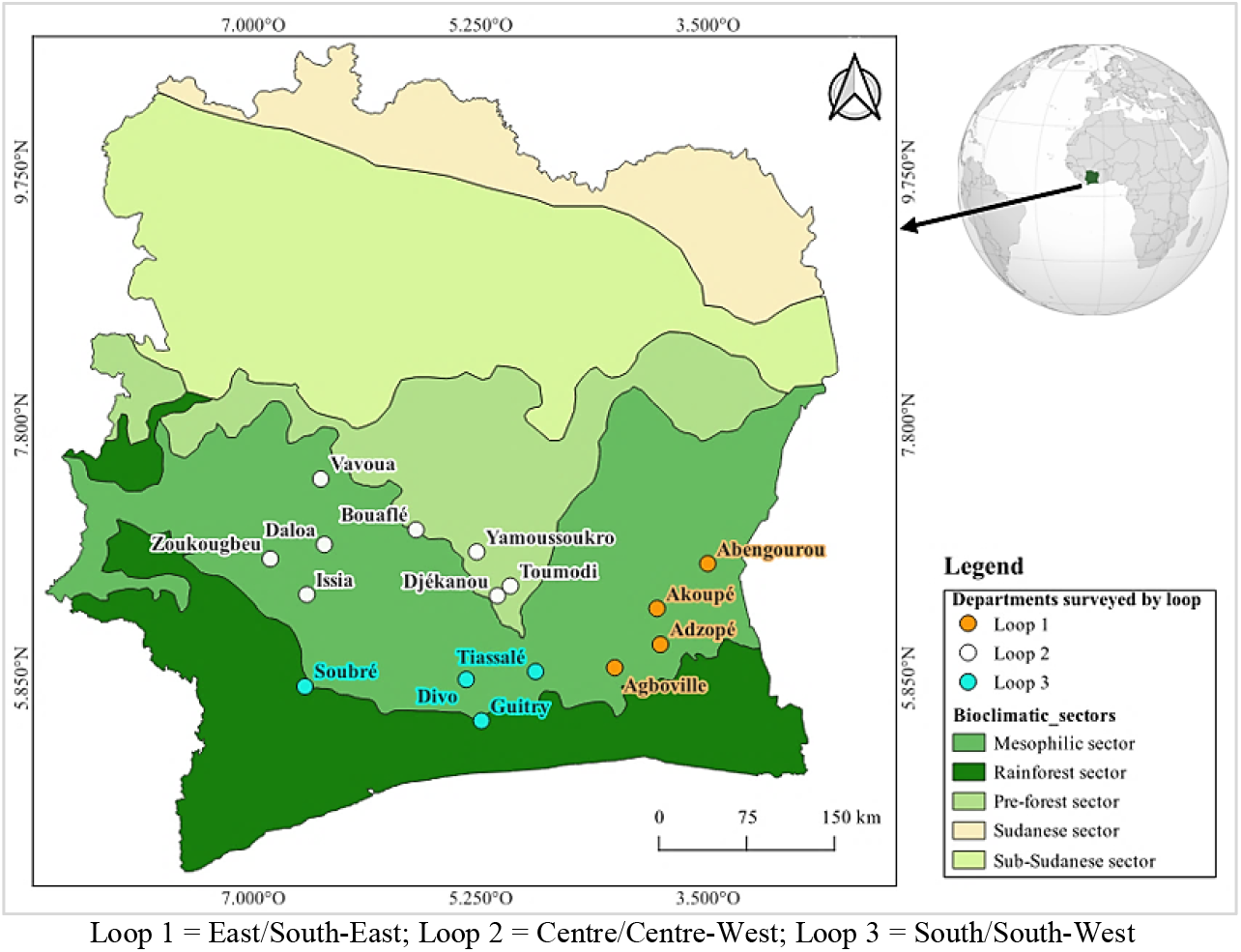
Location of the 16 survey departments (coloured by production loop), grouping the 103 recorded survey sites (villages and hamlets; see Table 1), within the bioclimatic sectors of Côte d’Ivoire, from the Sudanian sector in the north to the dense-forest sector in the south.

**Table 1.**
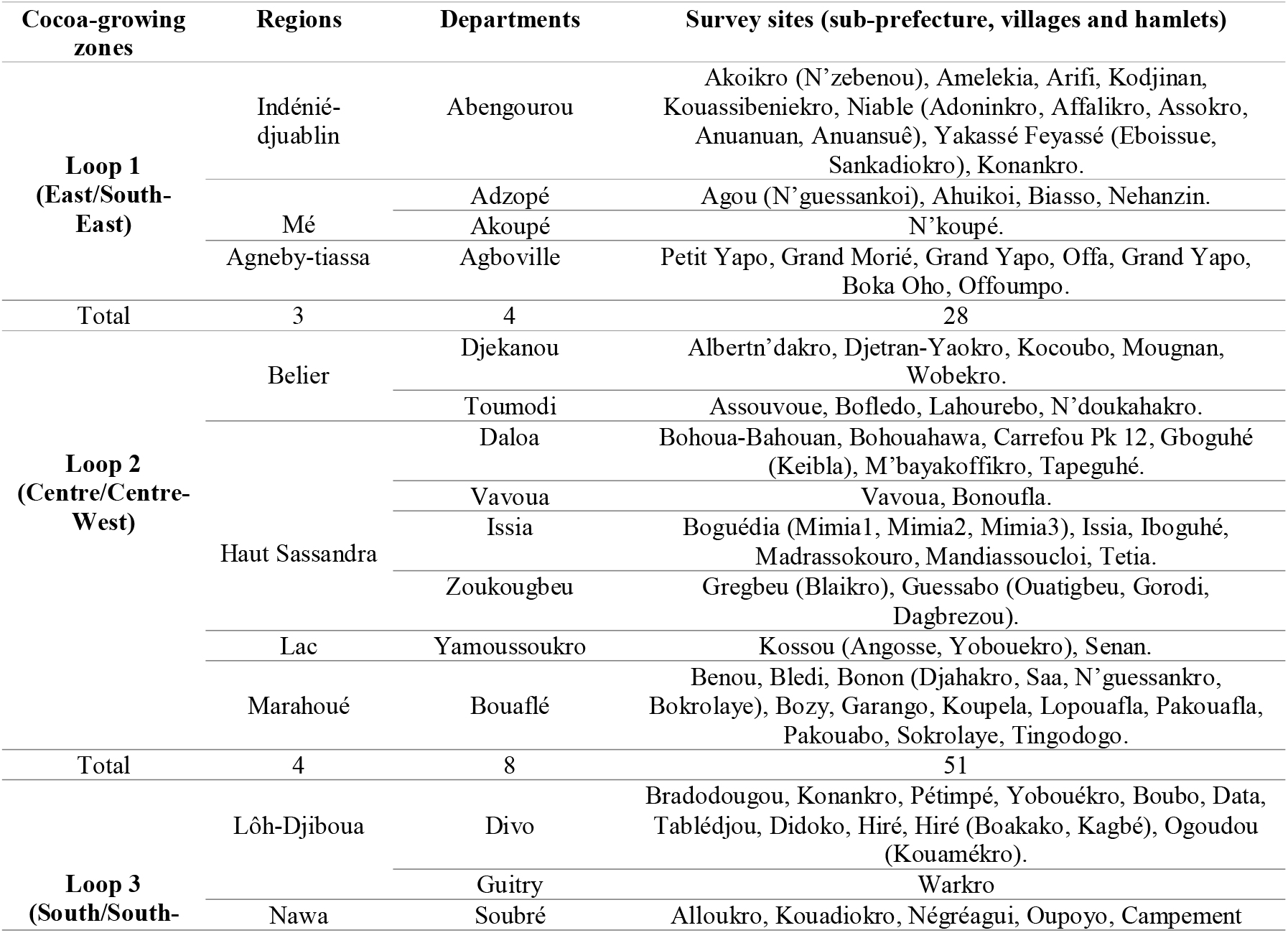
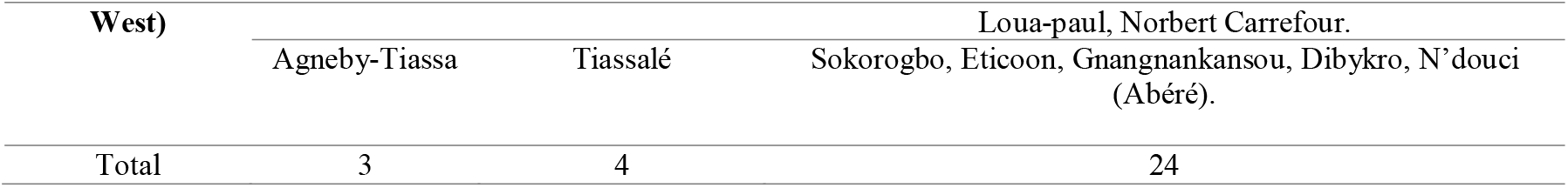
Presentation of survey sites by hamlet, village, sub-prefecture, department, region and loop.

**Table 2.** Descriptive statistics of the national sample (*n*=409, three loops)

| Variable | Mean / (%) | Standard deviation | Median | Min-Max |
| --- | --- | --- | --- | --- |
| Plantation age (years) | 24.2 | 15.8 | 20.0 | 1 - 83 |
| Plantation size (ha) | 3.6 | 4.3 | 2.5 | 0.2 - 46 |
| Yield (kg/ha/year) | 306.6 | 276.2 | 225.0 | 1.5 - 1837.5 |
| Farmer age (years) | 48.5 | 13.7 | 47 | 19 - 100 |
| Planted origin of shade trees | 82.9% | - | - | 339/409 |
| Aligned arrangement | 6.4% | - | - | 26/409 |
| Technical extension (ANADER/CNRA/SATMACI) | 18.1% | - | - | 74/409 |
| Selected cocoa planting material | 27.4% | - | - | 112/409 |
| Improved planting material (planted trees) | 12.7% | - | - | 52/409 |

For binary variables (%), the standard deviation and median are not reported (indicated by «-» ) since they add no information beyond the mean: the standard deviation of a proportion $p$ is entirely determined by it ( $\sqrt{p(1-p)}$ ), and its median is trivially equal to 0 or 1. For these same variables, the Min-Max column reports the number of observations showing the category of interest relative to the total sample ( $n/N$ ), rather than a range of values .

### 2.2. Data origin and geographic scope

The field survey was conducted in three successive waves between 2013 and 2016, in collaboration with the « Forest and Environment » research programme of the National Centre for Agronomic Research (CNRA). A closed, semi-closed and multiple-choice questionnaire, structured into four sections (general information on the farmer, agronomic characteristics of the plantation, characteristics of associated tree species, and management and ownership of trees within the agroforestry system), was developed and then piloted on a sample of farmers in the three loops before its final administration. The first wave (2013) covered four departments of the East/South-East loop (Abengourou, Akoupé, Adzopé, Agboville; 116 plantations, 461.15 ha); the second wave (2014) covered four departments of the South/South-West loop (Soubré, Guitry, Divo, Tiassalé; 180 plantations, 630.82 ha); the third wave (2016) covered eight departments of the Centre/Centre-West loop (Issia, Daloa, Zoukougbeu, Vavoua, Bouaflé, Yamoussoukro, Djékanou, Toumodi; 178 plantations, 504.75 ha), for a total of 474 plantations covering 1596.72 ha across 103 villages and hamlets (Table 1).

Within each sub-prefecture of the targeted cocoa-growing zones, villages were selected from among the main cocoa-production areas, based on information provided by the National Agency for Rural Development Support (ANADER) and, occasionally, on earlier CNRA work (location of on-farm experimental plots). Within each selected village, farmers were randomly selected, mainly from among members of cooperative sections active in the department and participants in ANADER farmer field schools; additional individual farmers were recruited directly in villages where these institutional relays were not available. Interviews were conducted individually, at the farmer’s home and directly on the plantation, with the support of technicians from the CNRA Cocoa Agronomy Laboratory.

The present article specifically draws on the socio-demographic (age, ethnic origin, education level, cooperative membership), agronomic (plantation age and size, origin and arrangement of shade trees, planting material, technical extension) and yield sections of this same questionnaire, for the sub-sample of plantations with a usable yield value. The « AVERAGE-YIELD » field of the original survey form (Supplementary material: S1) records the production declared over three consecutive annual campaigns, in bags, tonnes or kilograms depending on how the farmer reported it. A systematic audit of this field across all 470 usable plantations in the « GENERAL INFORMATION ON COCOA AND TREES » sheet (whose extraction required a specific correction, since the source sheet duplicated column headers for each of the three loops, with a column offset affecting loops 2 and 3 prior to correction) shows that 409 plantations have a usable yield value, spread across the three production zones (98 in loop 1, 151 in loop 2, and 160 in loop 3): individual yield is therefore not confined to a single geographic zone. After excluding non-numeric values and merging with plantation age and declared size, the final sample comprises 409 plantations spread across 124 villages in the three loops.

### 2.3. Construction of management variables

Annual yield per hectare was calculated as the mean (not the sum) of the three declared annual campaigns, each converted into kilograms (1 bag ≈ 75 kg) and then divided by the declared plantation size.

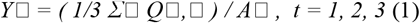

where *Y*□ is the annual yield per hectare of plantation *i* (kg·ha□^1^·year□^1^), *Q*□,□ is the declared production, converted into kg-equivalent, of plantation *i* for campaign *t* (*t* = 1, 2, 3, corresponding to successive campaign years), and *A*□ is the declared size of the plantation (ha).

Shade-tree management practice variables were decoded from the raw numeric codes of the survey form (Supplementary material: S1), which serves as the official codebook: origin of shade trees (residual/spontaneous/planted/other, multiple responses), arrangement (aligned/not aligned), planting material used for planted trees (wildlings/seeds/cuttings/marcots/grafts/other, recoded into « improved material » vs « basic material »), previous land use, mode of plantation acquisition, source of cocoa planting material (SATMACI/ANADER/CNRA-selected vs other), source of technical advice, and formal extension for the management of forest trees. Socio-demographic variables of the farmer (ethnicity, age, education level, cooperative membership) and behavioural variables (self-production of seedlings by the farmer, prior removal of forest trees, ownership of another plantation, number of perceived positive and negative effects of forest trees) were added in a second step. In total, 27 candidate variables were tested, including the production zone (loop) itself, systematically included as a covariate or interaction term to check the consistency of each association across the three zones rather than its significance in the pooled sample alone.

### 2.4. Variable excluded for insufficient data

The « DENSITY » (planting density, trees/ha) field remains insufficiently populated at the national scale (52 usable values out of 452 plantations, 11.5%) and was therefore excluded from the analyses, for lack of sufficient statistical power.

### 2.5. Resolution limit for a per-plantation diversity index

A floristic diversity index for shade trees at the scale of the individual plantation, which would have allowed a direct test of the diversity-yield trade-off hypothesis, could not be constructed uniformly across the whole sample: the floristic inventory of shade trees in loop 1 (« ASSOCIATED SPECIES INVENTORY » sheet) is available only at the level of three aggregated points (Abengourou, Adzopé, Agboville), rather than per individual plantation, unlike loops 2 and 3, where the inventory is available for 88 and 28 localities respectively. This asymmetry in data resolution prevents the construction of a per-plantation diversity index that is strictly comparable across the three zones, and motivated the reframing of the present study around management practices rather than an aggregated floristic diversity index.

### 2.6. Statistical strategy

The statistical analysis proceeds in four complementary steps, chosen for their respective strengths and limitations rather than to artificially converge on a single result, in order to assess the robustness of yield determinants across contrasting specifications:

✓ (i) Univariate screening: each candidate variable (among the 27 candidate variables drawn from the socio-demographic, agronomic and shade-tree management sections) is tested individually against the logarithm of yield per hectare (a transformation chosen to stabilise a strongly right-skewed distribution), using Pearson correlation for continuous variables and ANOVA/Kruskal-Wallis for categorical variables;
✓ (ii) A mixed-effects model (random intercept by village) on a pre-selected subset of predictors, with the production zone (loop) as a fixed covariate, to account both for the hierarchical spatial structure of the data (409 plantations in 124 villages) and for possible heterogeneity between zones:

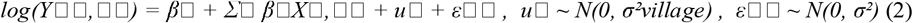

where *i* indexes plantations and *j* indexes villages, *X*□,□□ are the pre-selected covariates (age, size, production zone, planted origin, aligned arrangement, technical extension, education level), *u*□ is a village-level random intercept capturing unobserved inter-village heterogeneity, and *ε*□□ is the residual error term.

✓ (iii) An alternative production-function specification (logarithm of total annual yield as a function of the logarithm of plantation size and the same covariates, loop included), which avoids the instability of the kg/ha ratio for very small plots:

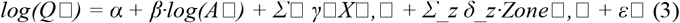

where *Q*□ is total annual production (kg), *A*□ is the plantation size (ha), *β* is the production elasticity of size (*β* = 1 corresponds to constant returns to scale), and *Zone*□,□ are the indicator variables for production zone (loop).

✓ (iv) *LASSO* (Least Absolute Shrinkage and Selection Operator) variable selection with calibrated penalty, and a random forest (Random Forest, 500 trees, maximum depth 4, minimum leaf size 4) with permutation importance, two machine-learning methods evaluated by 5-fold cross-validation on the full set of 27 variables (59 categories after encoding, loop included):

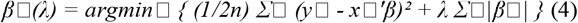

where λ is the *L1* penalty parameter (the penalty applies to the *L1* norm of the coefficients, i.e. the sum of their absolute values (*Σ*□|*β*□|), as opposed to an *L2* « sum-of-squares » penalty as in ridge regression), selected by 5-fold cross-validation to minimise the out-of-sample mean squared error; predictors retained with a non-zero *β*□□ are the variables selected by *LASSO* reported in the Results. This *L1*-penalisation selection approach is now well established in agronomic yield modelling (Shafiee et al., 2021). The random forest is parameterised with 500 trees, a maximum depth of 4, and a minimum leaf size of 4; its use, likewise validated by 5-fold cross-validation, is now documented for agricultural yield prediction (Jeong et al., 2016). Variable importance in the random forest (*y*) is measured by permutation: for variable *k*, it corresponds to the mean increase in out-of-sample prediction error when the values of *X*□ are randomly permuted across observations, with all other variables held fixed, over 50 repetitions.

For each of the methods above evaluated by cross-validation (mixed-effects model, production function, *LASSO*, and random forest), out-of-sample predictive capacity is summarised by the same indicator, the cross-validated *R*^*2*^:

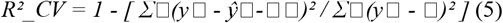

where *ŷ*□*-*□□ is the out-of-fold prediction for observation *i* under 5-fold cross-validation and □ is the sample mean of *y*; an *R*^*2*^_*CV* ≤ 0 indicates a model with no real predictive value beyond the sample mean. The out-of-sample predictive power of all models is reported using this cross-validated *R*^*2*^, alongside in-sample fit, throughout the Results.

For each emerging determinant, a systematic robustness check was conducted: (a) an interaction test with loop, to verify that the effect does not differ statistically from one zone to another; (b) replication of the test within each loop taken separately; (c) exclusion of extreme yield values (2.5th and 97.5th percentiles) to verify that the effect is not driven by a handful of influential observations. Out-of-sample predictive power (cross-validated *R*^*2*^) is systematically reported alongside in-sample fit, in line with good predictive-modelling practice: an in-sample *R*^*2*^ can be misleadingly high even in the absence of any generalisable signal, whereas a negative or null cross-validated *R*^*2*^ indicates that a model predicts no better than a simple mean. The nominal significance threshold adopted is α = 0.05, but the analysis explicitly accounts for the multiple-comparisons problem (27 variables tested) when interpreting univariate results. All analyses were conducted in Python 3.11 (pandas, numpy, scipy, statsmodels, scikit-learn 1.8).

## 3. Results

### 3.1. Sample overview

The final sample comprises 409 plantations (124 villages, three loops: 98 plantations in loop 1, 151 in loop 2, 160 in loop 3), with a mean age of 24.2 years (standard deviation 15.8), a mean size of 3.6 ha (median 2.5 ha, range 0.2-46 ha), and a mean yield of 306.6 kg/ha/year (median 225.0 kg/ha/year, standard deviation 276.2), with a strongly right-skewed distribution (Fig. 2A), justifying the logarithmic transformation adopted for modelling. The large majority of plantations (339/409, 82.9%) have shade trees of deliberately planted origin, while only a minority (26/409, 6.4%) show an aligned arrangement of shade trees. Although small, this proportion is, this time, sufficient to allow a statistical test of its effect (unlike an analysis restricted to loop 1 alone, where this variable is nearly constant).

**Fig. 2.**
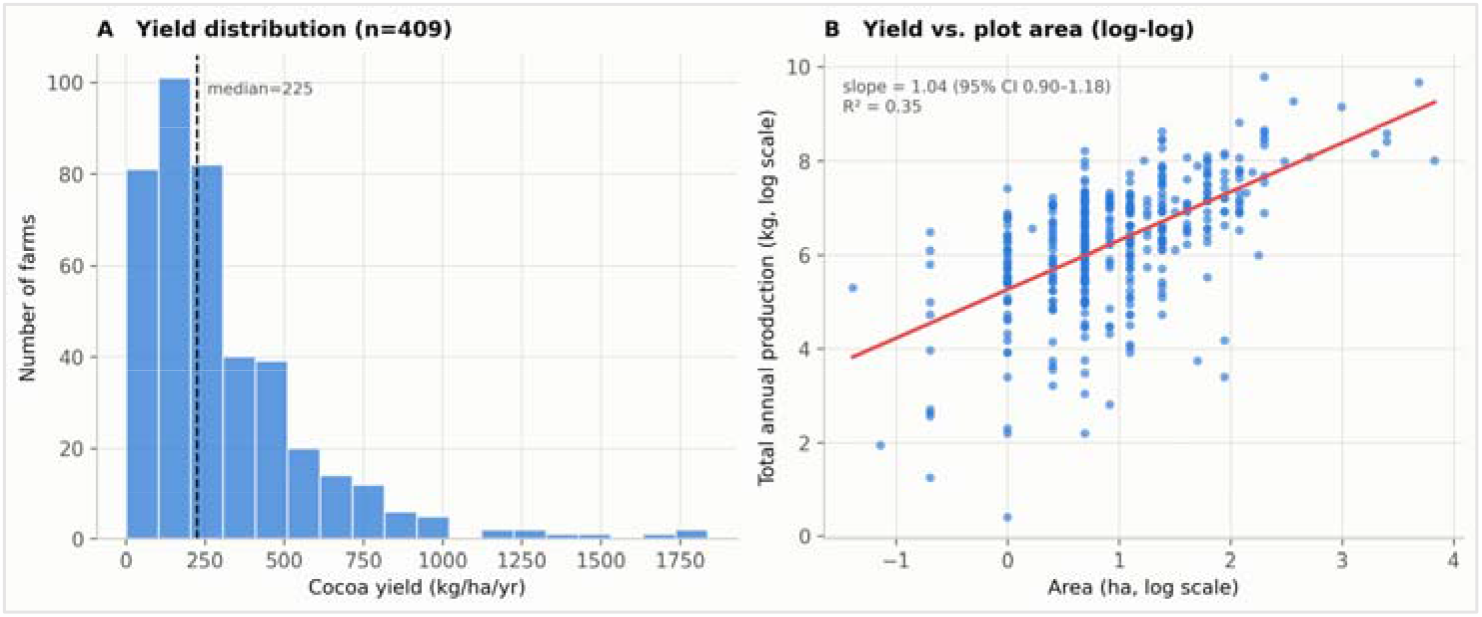
(A) Distribution of cocoa yield in the national sample (*n*=409, three loops) and (B) Total annual production as a function of plantation size (log-log transformation).

### 3.2. Univariate screening of practice variables

Univariate screening (Table 3) identifies seven variables crossing the nominal *p*<0.05 threshold (ANOVA and/or Kruskal-Wallis): the planted origin of shade trees (332 vs. 183 kg/ha/year in its absence, *p*<0.001), the presence of shade trees in the broad sense (311 vs. 218 kg/ha/year, *p*=0.028; a variable largely redundant with the previous one, since 388/409 plantations with shade trees include at least one of planted origin), agricultural technical extension (395 vs. 287 kg/ha/year, *p*=0.001), farmer education level (*p*=0.001, with the lowest yield among farmers with no schooling), the aligned arrangement of shade trees (456 vs. 296 kg/ha/year, *p*=0.020), the source of technical advice (*p*=0.019), and the planting material used for planted trees (*p*=0.027). None of the other 20 variables tested (plantation size, farmer age, residual/spontaneous origin of trees, previous land use, mode of acquisition, selected cocoa planting material, ethnicity, cooperative membership, self-reported behaviours) reaches this threshold, including the production zone (loop) itself (*p*=0.090 to 0.496 depending on the test), showing that mean yield differences between zones are not, by themselves, statistically significant when considered in isolation.

**Table 3.** Univariate screening of the 27 candidate variables (log-yield), ranked by increasing *p* value (Kruskal-Wallis test or Pearson correlation depending on variable type)

| Variables | Type | $n$ groups / $r$ | $p$ (Kruskal or Pearson) | Direction of effect |
| --- | --- | --- | --- | --- |
| Planted origin of shade trees | binary | 2 | <0.001 | Yes → higher yield |
| Technical extension | binary | 2 | 0.001 | Yes → higher yield |
| Education level | categorical | 4 | 0.001 | Primary/secondary → higher yield |
| Source of technical advice | categorical | 6 | 0.019 | Certification programme → higher yield |
| Aligned arrangement | binary | 2 | 0.020 | Yes → higher yield |
| Planting material (planted trees) | categorical | 4 | 0.027 | Improved → higher yield |
| Presence of shade trees | binary | 2 | 0.028 | Yes → higher yield (redundant with planted origin) |
| Production zone (loop) | categorical | 3 | 0.090-0.496 | n.s. |
| Aligned arrangement × loop (interaction) | test | - | 0.224 | n.s. (consistent effect across zones) |
| Plantation age | continuous | $r = +0.078$ | 0.114 | n.s. |
| All other variables (17) | various | - | > 0.15 | n.s. |

With 27 variables tested at the α=0.05 threshold, about one to two variables are expected to be significant by chance alone: the seven associations detected therefore clearly exceed this expected rate. A conventional univariate screening cannot, however, on its own rule out a chance association: each association requires independent verification before interpretation, in particular joint adjustment for confounding factors, a sensitivity assessment to extreme values, and a test of its out-of-sample predictive capacity. Fig. 3 illustrates, for descriptive purposes, the yield contrast observed for four of these variables: the planted origin of shade trees, technical extension, aligned arrangement, and farmer education level.

**Fig. 3.**
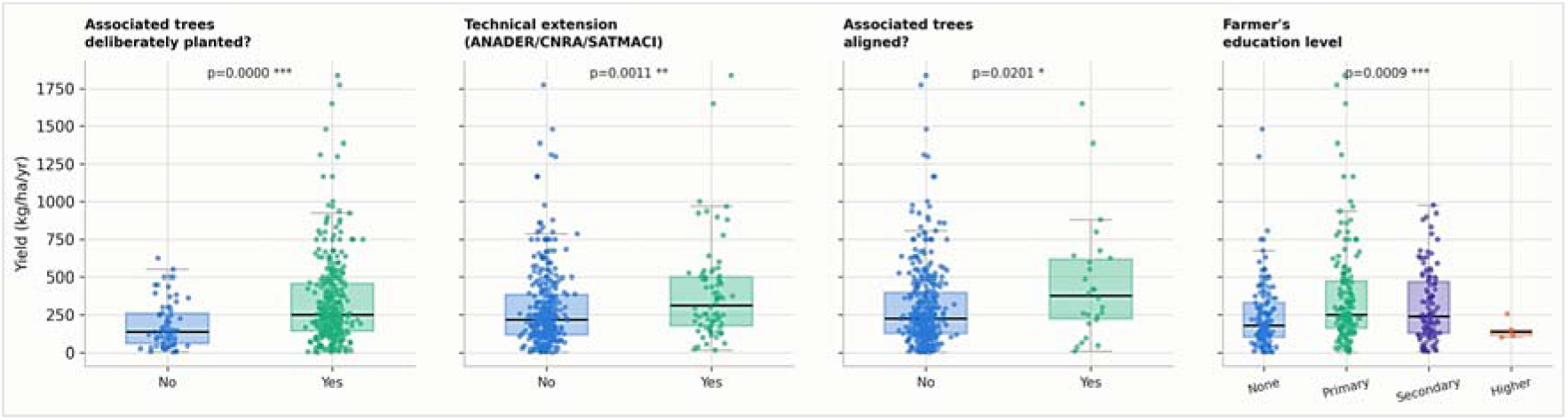
Yield by the four most robust determinants identified (national sample, *n*=409): deliberately planted origin of shade trees, agricultural technical extension, alignment of shade trees, and farmer education level.

### 3.3. Mixed model and production function

A multivariate model combining age, plantation size, production zone (loop), the planted origin of shade trees, aligned arrangement, technical extension, and farmer education level (*n*=409) achieves an in-sample *R*^*2*^ of 0.108 (*F*=4.82; *p*<0.0001). The planted origin of shade trees is the most robust predictor (with a coefficient of +0.52 on the log scale, corresponding to a yield multiplier of about 1.68, all else equal, *p*<0.001), followed by primary (+0.40, *p*=0.001) and secondary (+0.27, *p*=0.041) education relative to no schooling, and by technical extension (+0.31, *p*=0.021). Aligned arrangement remains positive but becomes marginal once adjusted for these other factors (+0.35, *p*=0.094; 26/409 aligned plantations). Production-zone coefficients are not significant (*p*=0.53 and 0.91 for loops 2 and 3 relative to loop 1), confirming that these associations are not an artefact of raw differences between zones.

The production-function specification (logarithm of total annual yield as a function of the logarithm of plantation size and the same covariates, Fig. 2B) confirms the consistency of these results (*R*^*2*^=0.396): the estimated slope for size is 1.01 (*95% CI* = 0.86-1.17), statistically compatible with a unitary elasticity, i.e. constant returns to scale (yield per hectare does not vary systematically with farm size). Technical extension (*p*=0.006) and education level (*p*<0.001 and *p*=0.012) remain significant in this alternative specification. A mixed-effects model with a random intercept by village also confirms the effect of technical extension (*p*=0.002) once the hierarchical spatial structure is accounted for.

### 3.4. LASSO and random-forest variable selection: a decisive robustness test

*LASSO* variable selection on the full set of 59 categories (27 variables, loop included) retains 6 non-zero coefficients: the planted origin of shade trees (the largest in absolute value), technical extension, primary education level, « Baoulé » ethnicity, a certification programme as a source of advice, and a residual planting-material category. The decisive test is that of out-of-sample predictive power: the 5-fold cross-validated *R*^*2*^ of this *LASSO* model is +0.054 on average (range +0.032 to +0.076 across folds, positive in all five folds), and that of a random forest on the same set of variables is +0.058 on average (positive in four folds out of five) (Fig. 4, panel B). A more restricted multivariate model, limited to the most robust variables (age, size, loop, education, planted origin, aligned arrangement, technical extension), achieves a slightly higher cross-validated *R*^*2*^ (+0.071 on average), and the planted origin of shade trees taken in isolation alone achieves a cross-validated *R*^*2*^ of +0.048 (positive in all five folds). This last variable is the simplest and most robust of the whole set tested.

**Fig. 4.**
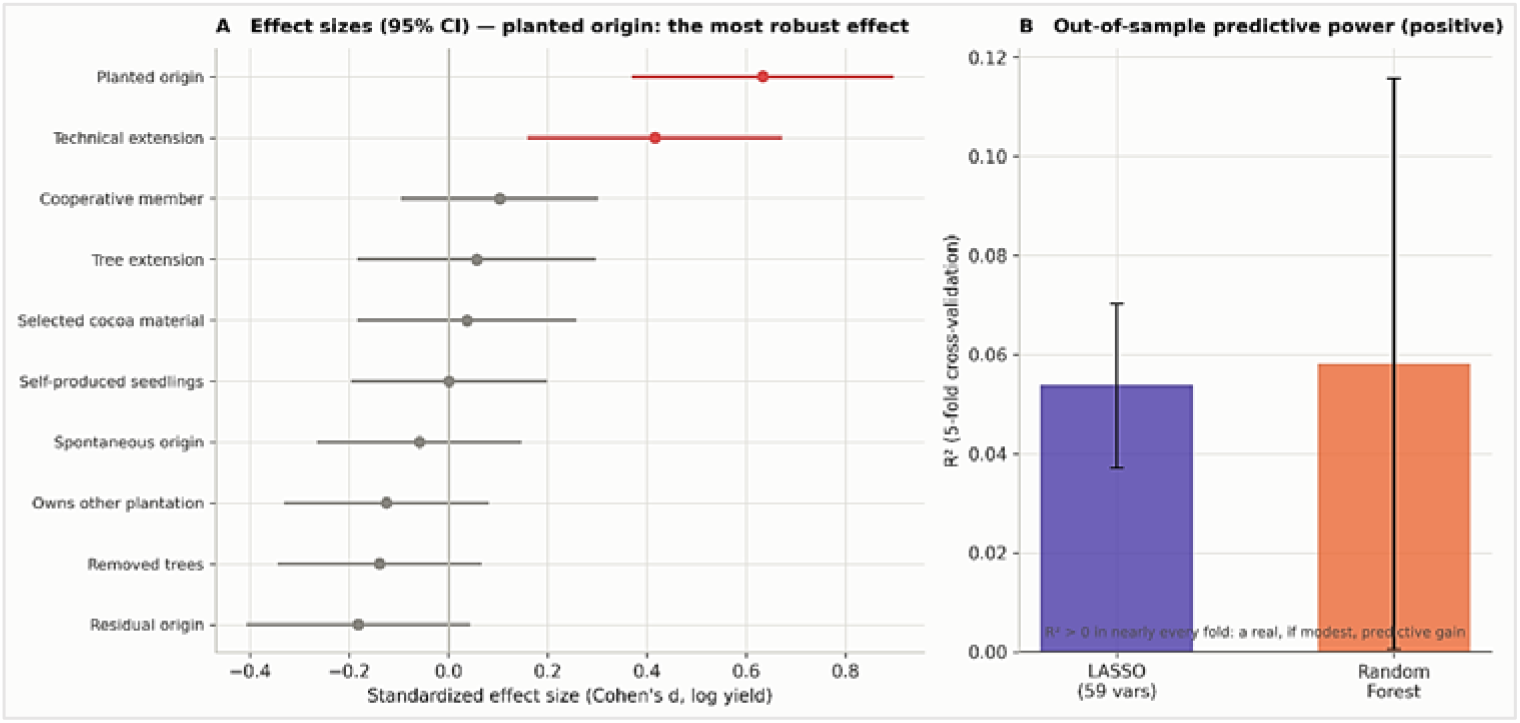
(A) Standardised effect sizes (Cohen’s *d*) and 95% confidence intervals for ten binary variables tested against the logarithm of yield; (B) Out-of-sample predictive power (5-fold cross-validated *R*^*2*^, positive) of a *LASSO* model and a random forest on the full set of 27 variables (59 categories).

Three robustness checks confirm that the effect of the planted origin of shade trees is not an artefact. First, it withstands the exclusion of extreme yield values (2.5th and 97.5th percentiles; *p*=0.0006 on the trimmed sample, *n*=387, versus *p*<0.001 on the full sample). Second, the interaction test with production zone is not significant (Type *II* ANOVA, *p*=0.224), and the effect is in the same direction within each of the three loops taken separately (loop 1: 234 vs. 320 kg/ha/year, Mann-Whitney test with *p*=0.373, not significant for lack of power with only 13 plantations without planted origin; loop 2: 175 vs. 389 kg/ha/year, *p*<0.001; loop 3: 153 vs. 300 kg/ha/year, *p*=0.012). Third, this effect is not confounded with the presence of shade trees in the broad sense: of the 409 plantations, only 21 have no shade trees at all, and almost all plantations with shade trees include at least one of planted origin (367/388), which explains the strong correlation between the two variables but confirms that planted origin captures a specific signal rather than a simple presence/absence contrast.

### 3.5. Visual synthesis of coefficients and LASSO / random-forest comparison

Table 4 gathers the main coefficients from the mixed model and the production function, giving direct access to the numerical values beyond the narrative description above. Fig. 5 (panel A) visualises the production-function coefficients as a forest plot: the 95% confidence interval clearly excludes zero for the logarithm of size, primary and secondary education, and technical extension, whereas it broadly includes zero for all the other covariates tested (plantation age, cooperative membership, planting-material origin, production zone), visually confirming the significance pattern presented above.

**Table 4.** Main coefficients of the mixed model (random intercept by village) and the production function (logarithmic scale)

| Model | Variable | Coefficient | *p* | Interpretation |
| --- | --- | --- | --- | --- |
| Production function | log(Size) | 1.014 | <0.001 | Elasticity $\approx 1$ (near-constant returns to scale) |
| Production function | Primary education | +0.524 | <0.001 | Higher total production |
| Production function | Secondary education | +0.359 | 0.012 | Higher total production |
| Production function | Higher education | +0.058 | 0.905 | n.s. ($n=5$, very small sample) |
| Production function | Technical extension | +0.390 | 0.006 | Higher total production |
| Production function | Cooperative membership | +0.055 | 0.620 | n.s. |
| Production function | Zone (loop 2 / loop 3 vs loop 1) | -0.151 / +0.017 | 0.311 / 0.907 | n.s. for both categories |
| Mixed model (village) | Technical extension | +0.433 | 0.002 | Higher yield/ha, village structure accounted for |
| Mixed model (village) | Spontaneous origin of trees | -0.185 | 0.112 | n.s. (negative trend) |
| Mixed model (village) | Size (ha) | -0.006 | 0.651 | n.s. |
| Mixed model (village) | Plantation age | +0.006 | 0.103 | n.s. (borderline) |
| Mixed model (village) | Inter-village variance | 0.171 | - | Moderate residual heterogeneity between villages |

Production function: $n=393$ , $R^2=0.396$ ; Mixed model: $n=409$ , 124 villages, *REML* method; n.s. = not significant at the $\alpha=0.05$ threshold

**Fig. 5.**
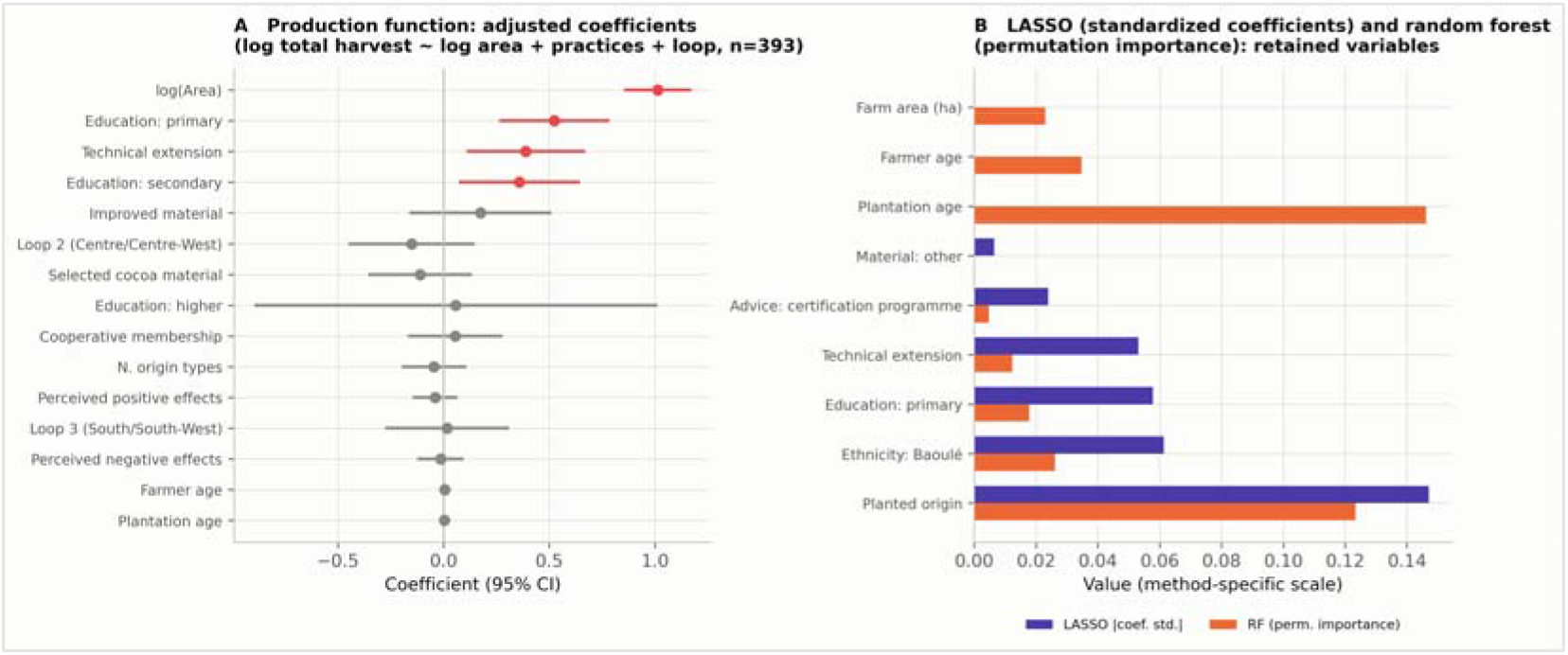
(A) Production-function coefficients (logarithmic scale) with 95% confidence intervals; (B) Comparison of the variables retained by *LASSO* selection (absolute value of the standardised coefficient) and by random-forest permutation importance, for the union of the most important variables according to each method.

Panel B of Fig. 5 compares the variables retained by *LASSO* selection (absolute value of the standardised coefficient) and by random-forest permutation importance, for the union of the most important variables according to each of the two methods. Both methods converge on the planted origin of shade trees as the dominant variable, but diverge notably on plantation age: *LASSO* assigns it a near-zero coefficient (a variable discarded by *L1* penalisation in favour of predictors more strongly correlated with yield), whereas the random forest assigns it the highest permutation importance among all the variables tested. This methodological divergence, rather than a contradiction, illustrates a well-documented difference in behaviour between penalised regression and tree-based ensemble methods when faced with correlated predictors or non-linear relationships: it cautions against over-interpreting the absence of a variable from a *LASSO* model as proof of its lack of relevance.

## 4. Discussion

### 4.1. The planted origin of shade trees: a robust determinant of yield

The central result of this study is that the deliberately planted origin of shade trees associated with cocoa is the strongest and most stable determinant of individual yield, with a mean gap of 332 versus 183 kg/ha/year between plantations with planted trees and those with residual or spontaneous-origin trees. This effect confirms hypothesis *H1* and withstands a set of convergent checks rather than a single favourable specification: it remains significant after simultaneous adjustment for age, size, production zone, technical extension, and education level; it is not driven by a narrow subset of extreme plantations (*p*=0.0006 on the trimmed sample of the 2.5th-97.5th percentiles, versus *p*<0.001 on the full sample); it holds, in the same direction, within each of the three production zones taken separately rather than in only one of them; and, taken in isolation, it retains a positive out-of-sample predictive power (cross-validated *R*^*2*^ ≈ 0.048), comparable to that of the full *LASSO* model built on all 27 variables (*R*^*2*^ ≈ 0.054). A determinant this simple to measure and to communicate to farmers deserves, on this basis alone, to be regarded as a priority for agroforestry extension rather than as merely one statistical result among others.

### 4.2. A planted origin reflects a deliberate farmer choice, not a mere by-product of natural regeneration

The most parsimonious interpretation of this effect is that planted origin captures an active investment by the farmer (species choice, planting effort, and subsequent maintenance) rather than a simple presence/absence contrast for shade trees: of the 409 plantations in the sample, only 21 have no shade trees at all, and almost all plantations with shade trees (367/388) include at least one of planted origin. This interpretation finds direct support in the Ivorian context: across deforested cocoa landscapes of Côte d’Ivoire, farmers’ choice of species actually planted reflects deliberate trade-offs between use value, market value, and compatibility with cocoa (Atangana et al., 2021), indicating an active, selective decision rather than a mere by-product of natural regeneration. At the continental scale, the presence of trees on the farm remains a significant and widespread economic choice among smallholders (Miller et al., 2017), and recent work in Nigeria confirms that it is farmers’ perceptions of the benefits derived from planted trees, more than their mere availability, that drive the decision to plant (Olatujoye et al., 2024). A complementary structural determinant, not directly measured here, could reinforce this interpretation: land tenure security is a long-standing and robust driver of tree-planting incentives in Africa (Fortman, 1985; Mekonnen, 2009), suggesting that part of the effect attributed here to planted origin could also reflect, upstream, greater tenure security among the farmers concerned.

### 4.3. Alignment of shade trees: partial convergence with recent Ivorian work

A very recent, independent study on a distinct sample of Ivorian plantations (Yéo et al., 2026) identified the alignment of planted trees as the strongest determinant of cocoa yield (+51%), followed by mineral fertilisation (+29%) and certified planting material (+23%). Our own sample recovers a univariate signal in the same direction and of comparable magnitude for alignment (456 vs. 296 kg/ha/year, i.e. +54%), but this effect becomes statistically marginal once jointly adjusted for planted origin, technical extension, and education (+0.35 on the log scale, *p*=0.094), and it concerns a narrow subgroup of plantations (26/409). This divergence is not necessarily contradictory: alignment and planted origin are two related but distinct dimensions of management intentionality (a tree can be planted without being aligned, and vice versa), and it is plausible that the sample of Yéo et al. (2026), constituted differently, captures a context in which alignment is a more discriminating marker of intensive management. This finding underlines the value of clarifying, in future work, which of these two dimensions (the origin or the arrangement of shade trees) carries the more robust causal signal, rather than treating them as interchangeable.

### 4.4. Agricultural human capital: technical extension and farmer education as robust, second-tier determinants

Agricultural technical extension (access to ANADER, CNRA, SATMACI advice, or a certification programme) and farmer education level emerge as robust, second-tier determinants, partially confirming H2: both remain significant in the mixed model and in the production function, and are retained by *LASSO* selection. This finding is consistent with a recent meta-analysis spanning a wide range of agricultural contexts, which concludes that access to technical extension has a positive average effect on productivity, while highlighting substantial heterogeneity depending on the quality and nature of the services provided (Ogundari, 2022). In the same vein, recent panel data from Ghana confirm a comparable positive effect of extension on farm-household welfare (Aremu et al., 2025), while in South Africa, more broadly defined human capital likewise contributes positively to the productivity of family farms (Baiyegunhi, 2024). An important caveat regarding causal interpretation should nonetheless be noted: farmers who seek out technical extension, or who have attained a higher level of education, may systematically differ, along dimensions not observed here (motivation, access to credit, social capital), from farmers who do not. This constitutes a selection bias that a purely observational design such as this one cannot fully rule out.

### 4.5. Farm size and cooperative membership are not robust determinants of yield

In line with hypothesis H3, neither plantation size nor cooperative membership emerges as a robust determinant of yield once management practices and the farmer’s human capital are accounted for: neither variable reaches the significance threshold at univariate screening, and the production function reveals an elasticity of total yield to size that is statistically indistinguishable from 1 (estimated slope 1.01, 95%CI = 0.86-1.17), meaning that yield per hectare is broadly independent of farm size in our sample. This finding of near-constant returns to scale echoes the conclusion of Ali and Deininger (2015) for Rwandan agriculture. More recent re-examinations of this relationship, covering a wider range of farm sizes, reach similar conclusions in Kenya (Muyanga and Jayne, 2019) and Nigeria (Omotilewa et al., 2021), while a study focusing specifically on Ghanaian cocoa concludes that spatial and management determinants, rather than size itself, explain most of the productivity variation between farms (Bentum et al., 2026); a result that directly echoes the one obtained here. Regarding cooperative membership, a recent Ethiopian study of maize farmers finds equally mixed results depending on the specification used, reinforcing the view that its effect on productivity depends heavily on institutional context and estimation method (Geffersa, 2024); in the specific context of compliance with new European traceability requirements, a study conducted in western Côte d’Ivoire further suggests that cooperative membership mainly promotes documentary compliance rather than productive performance as such (Moluh Njoya et al., 2025), which could explain why a simple membership indicator, such as the one used here, does not capture a direct effect on yield.

### 4.6. An instructive divergence between LASSO and random forest

*LASSO* variable selection and random-forest permutation importance converge on the planted origin of shade trees as the dominant variable, which strengthens confidence in this central result beyond a single family of methods. The two approaches diverge markedly, however, on plantation age, to which *LASSO* assigns a near-zero coefficient while the random forest assigns it the highest importance among all variables tested; a divergence that is not specific to our dataset: *L1*-penalised regression tends to retain only one representative from a group of correlated predictors, or to ignore non-monotonic relationships, whereas tree-based ensemble methods remain sensitive to both (Shafiee et al., 2021, for a comparable discussion in agronomic yield modelling). This divergence cautions against too quickly concluding that a variable discarded by *LASSO* lacks agronomic relevance, and argues for the systematic use of several families of methods rather than a single one, as was done here.

### 4.7. Limitations

Several limitations should be noted. First, yield is a retrospective farmer self-report covering three agricultural campaigns, subject to potentially non-negligible measurement error, although nothing indicates that this error is correlated with the determinants studied. Second, the three survey waves each correspond to a distinct calendar year (2013 for loop 1, 2014 for loop 3, 2016 for loop 2): production zone and survey year are therefore partially confounded, so that the absence of a detected zone effect cannot be interpreted as the strict absence of any geographic heterogeneity independent of the survey calendar. Third, the design is observational and cross-sectional: it cannot rule out a selection bias whereby farmers who deliberately plant shade trees, seek out technical extension, or pursue their education systematically differ, along unobserved dimensions, from farmers who do not. Fourth, as noted in the Methods, the heterogeneous resolution of the shade-tree floristic inventory did not allow the construction of a per-plantation diversity index that is strictly comparable across the three zones, leaving untested the diversity-yield trade-off hypothesis that is nonetheless central to the cocoa agroforestry literature (Clough et al., 2011; Tscharntke et al., 2011).

## 5. Conclusion

This study establishes, on a nationwide sample of 409 plantations covering, for the first time, the three main cocoa-production zones of Côte d’Ivoire, that the deliberately planted origin of shade trees associated with cocoa is the strongest and most stable determinant of individual yield; a result that withstands four independent robustness checks (multivariate adjustment, exclusion of extreme values, consistency across the three production zones, and positive out-of-sample predictive power) rather than resting on a single favourable specification. Agricultural technical extension and farmer education level constitute a second group of robust determinants, albeit of more modest magnitude, while farm size and cooperative membership do not emerge as independent determinants of yield, confirming the competing hypotheses formulated a priori. Methodologically, the convergence of four complementary approaches (univariate screening, mixed model, production function, and machine learning « *LASSO* and random forest ») systematically evaluated by cross-validation rather than by in-sample fit alone illustrates the value of a multi-method robustness approach for detecting agronomic determinants that are modest in magnitude but real, in noisy survey data; an approach transposable to other agroforestry contexts. These results argue for a more systematic integration of shade-tree origin (alongside technical extension and education) into Ivorian agroforestry extension programmes, while calling for a dedicated experimental or longitudinal design to establish the causality of this effect and refine its measurement at the plantation scale.

## Author Contributions

BIA. Developed the methodology, collected survey data, analysed the data and wrote the paper. AAA. Developed the methodology, collected survey data and supervised the work. AME, KKE and DSA supervised the work.

## Fundind

This work was funded by the National Centre for Agricultural Research of Côte d’Ivoire (*CNRA*).

## Declaration of competing interest

The author declare no competing interests.

## Acknowledgements

The authors would like to thank the rural communities in the surveyed areas for their hospitality, kindness and cooperation, and for making their plantations available to ensure the smooth implementation and execution of the activities carried out on their land during this period. The authors would also like to thank ANADER Côte d’Ivoire and SATMACI (*Société d’Assistance Technique pour la Modernisation de l’Agriculture en Côte d’Ivoire*) for its support, for making the cooperatives and cocoa farmers in the surveyed areas available, and for facilitating access to them. The authors would like to express their sincere thanks to Miss Houphouet Aya Diane Larissa for producing the map showing the distribution of the study’s sampling sites.

## Data availability

The datasets analysed as part of this study are available at: Adji, B. I., Assiri, A. A., Assi, M. E., & Kassin, K. E. (2026). Raw survey and floristic inventory data on cocoa-based agroforestry systems in Côte d’Ivoire (three cocoa-growing “loops”, 2013-2016): anonymized version [Dataset]. Zenodo. https://doi.org/10.5281/zenodo.22016383

## Notes

https://zenodo.org/records/21435509

### Competing Interest Statement

The authors have declared no competing interest.

